# A Straightforward Approach for Living Biomembrane Printing onto Nanoparticle

**DOI:** 10.1101/2023.10.22.563496

**Authors:** Ryosuke Mizuta, Eisuke Kanao, Keigo Ukyo, Shin-ichi Sawada, Yasushi Ishihama, Yoshihiro Sasaki, Kazunari Akiyoshi

## Abstract

Biomembrane coating technologies have increasingly been pursued to grant natural dynamic bio-interfaces onto synthetic nanomaterials. Herein, we report a one-step method to coat “living” biomembrane on nanoparticle surfaces in a non-destructive manner. In our method, nanoparticles were efficiently coated with cell membranes without losing the structural integrity by mechanically facilitating the passage of nanoparticles to a concentration layer of living cells with simple centrifugation. This was similar to the exosome-releasing process via endocytosis and exocytosis. The biomembrane originating from living Raw264.7 cells was coated onto the silica nanoparticle prepared by our method, and proteome profiling with nanoflow liquid chromatography-tandem mass spectrometry demonstrated that it was constructed with proteins derived from the membranous component. This proteome profile was not observed in silica nanoparticles prepared with dead cells. Finally, the hybridized cell membrane effectively suppressed the phagocytic activity of Raw264.7 cells to silica nanoparticles and improved the uptake efficiency into cancer cells. We believe our simple and efficient method to coat living biomembranes should be useful in developing medical and pharmaceutical applications involving nanoparticles.

## Introduction

Nanotechnology has attracted great attention over the last decade due to its numerous biomedical applications, including targeted drug delivery systems (DDS), bioimaging, and cancer treatment^1, 2, 3^. A systematic understanding of nano−bio interaction is needed for the intelligent design of safe and effective engineered nanomaterials for biomedical applications because they are recognized as foreign and could be directed to the appropriate cellular elimination system by an immune defensive response^4^. In traditional chemical/physical approaches, such as PEG functionalization, morphology control, and lipid modification, efforts have been made to construct regulated nano-bio interfaces to sneak the nanomaterials into exclusive biological systems^5, 6, 7^. Researchers have aimed to design engineered nanomaterials with the ultimate goal of being ignored by all cells except the target cells; however, negative immunogenic responses and allergic reactions were unavoidable^8^. In addition, the need for complex chemical treatments and synthetic modifications has made the system overly complex and limited its universal applications in biomedicine. There is still significant room for designing an integrated biomimetic nano-interface.

Natural biomembrane nanoparticles, emerging as novel biotic nano-carriers that control physiological and pathological scenarios, have vital clues to improve the biointerfacing capabilities of engineered nanomaterials^9, 10^. With the lipid bilayers derived from donor cells, natural biomembrane nanoparticles have the targeting ability of specific surface proteins or glycans to accurately deliver encapsulated substances to recipient cells^11^. For example, exosomes are cell-secreted nanoparticles (generally with a size of 30–150 nm) and mediate local and systemic intercellular communication^12^.

Exosome uptake is controlled by interactions between exosome surface proteins and recipient cells, regulating immune responses, cancer metastasis sites, and other biological processes^13, 14^. In fact, the adhesion-associated molecules on the exosome surface, such as tetraspanins, integrins, and glycan, determine which cells receive exosomes^15, 16, 17^. Another example is enveloped viruses, which acquire membrane structures from host cells by coating themselves with host cell-derived plasma membranes or nuclear membranes during proliferation and budding from host cells^18^. The membrane proteins encoded by viral genes are expressed on these membranes, involving cell attachment and entry^19^. There are some reports that aim to hybridize such biological membrane nanoparticles with nanomaterials for applications in biomedicinal fields^20, 21^. However, the production of natural nanoparticles has relied on the biological process of living cells for production, and there have been unsolved problems such as low isolation efficiency, insufficient target specificity, compositional heterogeneity, and low drug loading efficiency^22, 23, 24, 25, 26^.

On the other hand, biomembrane coating technology, hybridizing natural biomembranes and synthetic nanoparticles, has garnered attention^27^. These methods have enabled us to artificially confer the multifaceted advantages derived from natural biomembrane vesicles, such as limited immunogenicity, improved circulation stability, unique targeting capabilities, and enhanced intracellular delivery efficiency, onto nanomaterials with unique functionalities^28, 29^. The technology has already been applied to different types of biomembranes, including red blood cells (RBCs), platelets, white blood cells, cancer cells, stem cells, and even bacteria^30, 31, 32, 33, 34, 35^. However, most of the methods were based on membrane components extracted by destroying cells^29, 36^. In these methods, the membrane components were disrupted by ultrasound or extrusion of cells, and the biomembrane hybridized nanoparticles were prepared by sonication of these membrane components in the presence of core particles. In fact, these conventional methods have pointed out the possibility that these conventional methods could lack the structural integrity of the cell membrane coating on the particle surface. Low coating efficiencies, less than 10%, were also a big problem of the conventional method due to applying mechanical energy to the extracted membrane components^37^. Therefore, a rational strategy has been highly desired to efficiently coat membranous components to the artificial nanoparticle surface without compromising the integrity of the biomembrane.

In our previous study, we showed that nanoparticles mechanically penetrating concentrated lipid layers can efficiently coat the particle surface with phospholipid bilayers^38^. Herein, we newly improved this technique by applying living cells themselves instead of lipid layers (Figure 1), for effectively coating “living” membranes onto nanoparticles. In particular, cells were concentrated at the interface of solutions of different densities, and silica nanoparticles were mechanically penetrated through the layers with simple centrifugation. This was similar to the exosome-releasing process via endocytosis and exocytosis. By optimizing the composition of the solution, cell viability could be maintained before and after penetration. Briefly, this method is extremely unique in effectively coating snapshots of the biomembranes using living cells as a source. Furthermore, the difference in proteome profiles of biomembrane hybridized silica nanoparticles prepared with living and dead cells was clearly demonstrated by nanoflow liquid chromatography-tandem mass spectrometry (LC-MS/MS); membranous proteins were predominantly enriched on the nanoparticles prepared with living cells. The simplicity, efficiency, and characteristic molecular composition derived from living cells suggest that our method could be a universal biomembrane coating platform for the biomedical application of a wide range of artificial nanomaterials.

**Figure 1.**
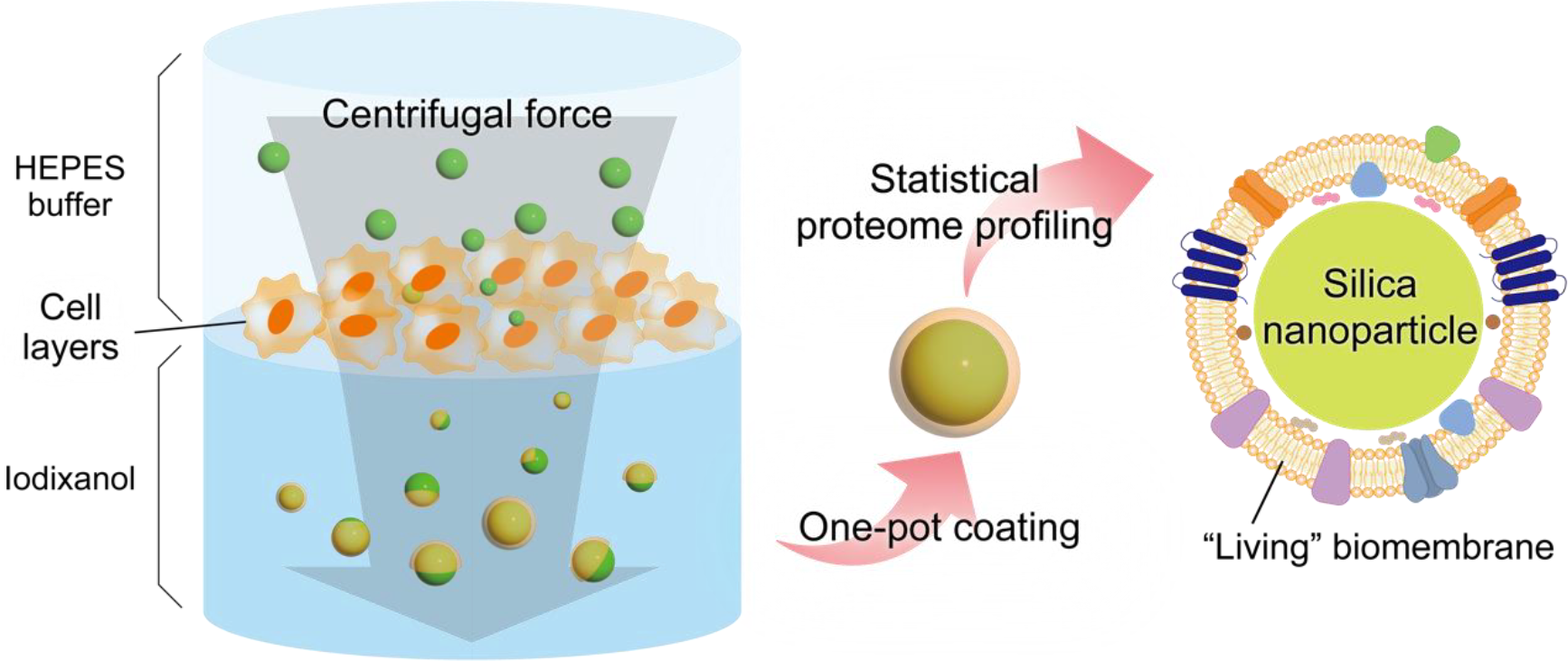
Schematic diagram of the biomembrane coating method. Nanoparticles were efficiently coated with cell membranes without losing the structural integrity by mechanically facilitating the passage of nanoparticles to a concentration layer of living cells with simple centrifugation.

## Results and discussion

### Optimization of conditions enables biomembrane coating using living cells

Inspired by the method of fractionating cells by density gradient centrifugation, we attempted to form “living” cell layers at the interface of solutions of different densities. By using living cell layers, the source cells could keep the integrity of the membrane without the complex process involving the extraction of biomembranes. Firstly, we investigated sucrose, which is commonly used in density gradient centrifugation. 60 wt% sucrose solution was suspended with Raw264.7 cells and stacked with HEPES buffer to form a two-phase system with different densities. Ultracentrifugation was then performed to confirm that cell layers did indeed form at the interface between these solutions. Fluorescently labeled silica nanoparticles with a particle size of 100 nm were added from the top of these cell layers, and the silica nanoparticles were penetrated by ultracentrifugation for 2 hours. The particle size measurement by DLS of the particles obtained after purification showed an increase in particle size of about 20 nm with penetration of the cell layers (Figure 2A). This suggested that cell membrane components may have been coated onto the surface of the silica nanoparticles by the penetration of the cell layers. The yield of silica nanoparticles calculated from the fluorescence intensity of FITC modified on silica nanoparticles was about 65%, indicating that silica nanoparticles can be obtained in relatively high yields. Furthermore, the ζ-potential measurement of the obtained particles in the HEPES buffer showed that the potential of the silica nanoparticles, which were largely negatively charged, was relaxed by penetration of the cell layers (Figure 2B). ζ-potential changes are widely used as evidence of cell membrane coating on the particles^43^. Here, the charge relaxation observed strongly suggested that the cell membrane has been complexed to the silica nanoparticle surface.

**Figure 2.**
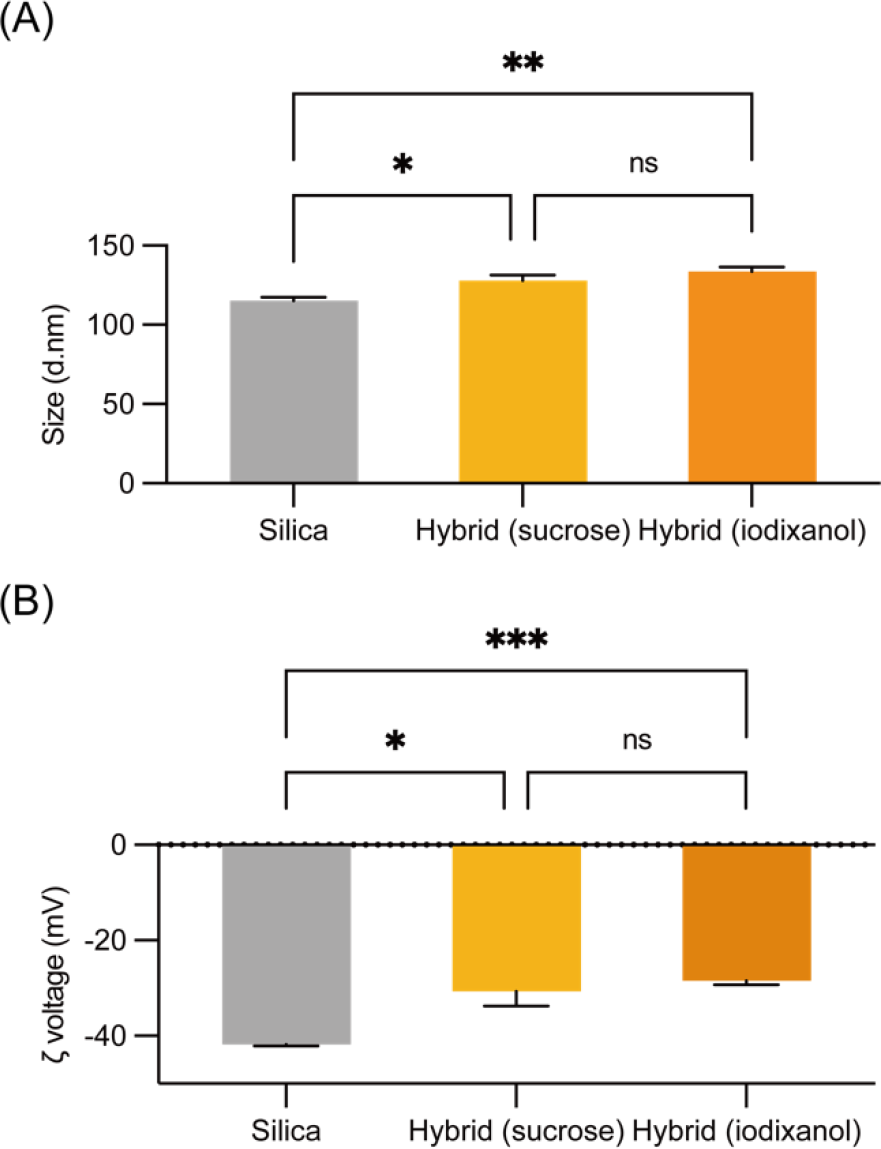
(A) Size change and (B) ζ-potential change of silica nanoparticles penetrating through the cell layers formed by each density gradient solution. “Hybrid” here refers to silica nanoparticles after penetration through the cell layer. All the experiments were repeated for three times (n = 3) and data were presented as mean ± s.d.

However, when cell viability after the penetration was measured by Toripan Blue staining, it was clear that almost all cells had lost their membrane integrity (Figure S1). This result was also confirmed when a similar manipulation was performed without the addition of silica nanoparticles. Thus, the method using sucrose was shown to be invasive to cells due to the high osmotic pressure and the longer centrifugation time required for silica nanoparticle penetration due to the increased solution viscosity. Therefore, we formed layers using a solution for density gradients that is less invasive to cells. We focused on Iodixanol, a versatile centrifugation medium that can be used to separate extracellular vesicles and cells^39, 40^. Iodixanol has a lower viscosity and osmotic pressure than sucrose, and thus, as we had expected, can reduce damage to cells during penetration. We therefore considered that 30 wt% Iodixanol could be used for the penetration. We performed cell layer formation and penetration of silica nanoparticles using 30 wt% Iodixanol solution, and most of the added silica nanoparticles were obtained in 10 minutes when Iodixanol solution was used for the lower layer. Measurements of particle size and ζ-potential showed an increase in particle size and relaxation of surface potential with penetration of the cell layers, and the biomembrane-hybridized nanoparticles were prepared similarly to those prepared with sucrose. The cell viability after the biomembrane coating operation was relatively high, around 80%, suggesting that membranes derived from living cells can be coated on the nanoparticle surface by optimizing the solution for the density gradient.

### Proteome profiling of biomembrane hybridized silica nanoparticles

Biomembrane coating efficiency is an important indicator in assessing the quality of biomembrane-coated particles. However, not many studies mentioned the coating efficiency. It has been also clear that conventional coating methods such as ultrasonic irradiation or extruder treatment in the presence of extracted membrane components and nanoparticles can mostly only partially coat the nanoparticles. Therefore, we investigated the biomembrane coating efficiency of particles prepared by our method. Specifically, we used the fluorescence quenching method of NBD^37^. We treated NBD-labeled silica nanoparticles with dithionite, a reducing agent that does not penetrate cell membranes, to irreversibly quench NBD fluorescence (Figure 3A). Initially, the addition of quenching agents to bare silica nanoparticles resulted in a marked decrease in particle-derived fluorescence. In contrast, for biomembrane hybridized particles prepared with cell layers formed from living cells using Iodixanol, 20% of the fluorescence before the addition of the quenching agent was still detected after the addition of the quenching agent. This clearly showed that 20% of the total particles were completely coated with the biomembrane. Since the complete coverage was less than 10% in the conventional method of hybrid particles, it was suggested that this method can efficiently coat the surface of nanoparticles with biomembrane. On the other hand, almost all fluorescence was quenched in the particles prepared with sucrose. This indicated that particles prepared with dead cells as a source have incomplete coverage. Therefore, it was emphasized that coating particles with living cells as a source is important to achieve higher coverage.

**Figure 3.**
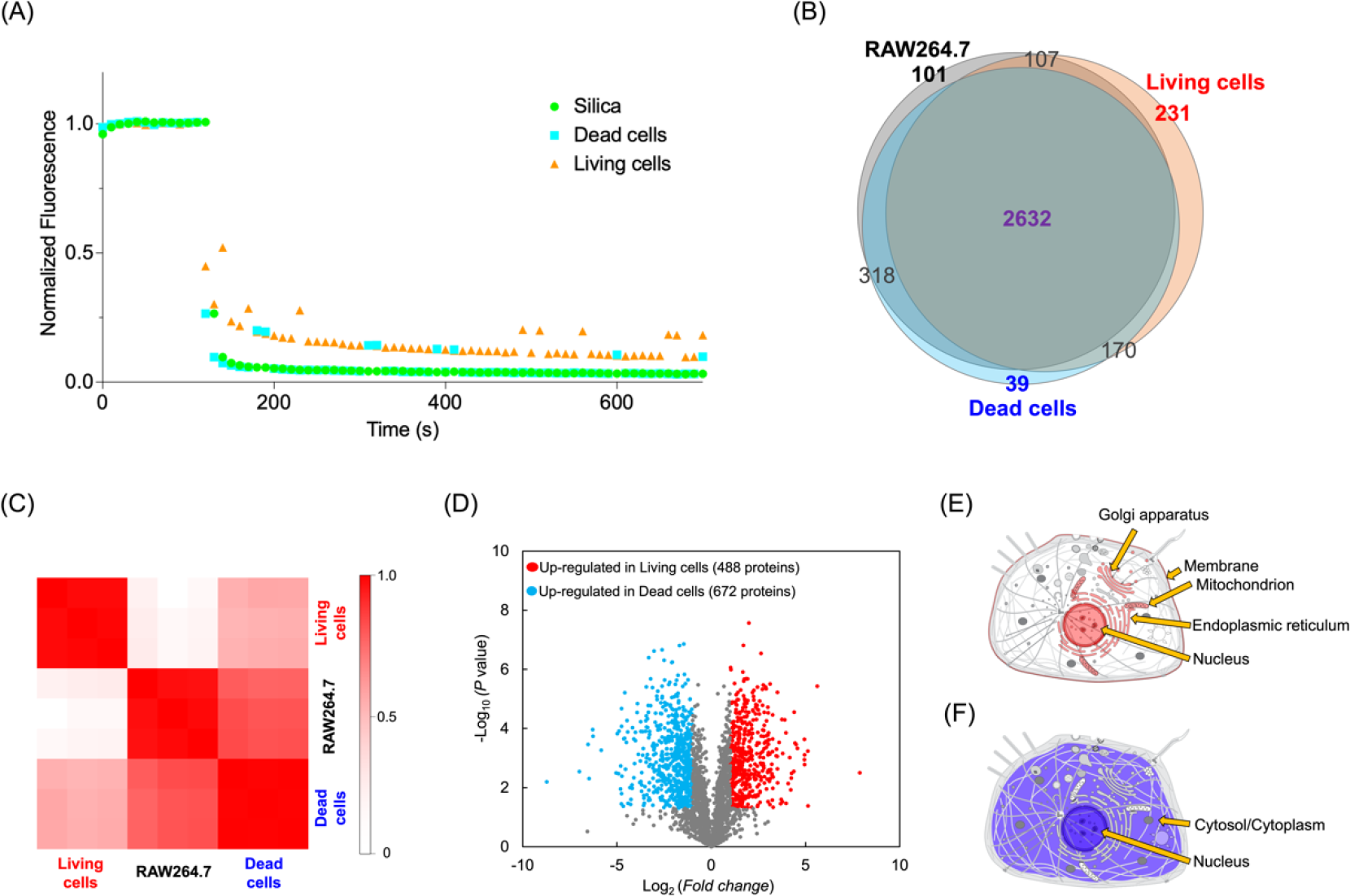
Characterization of biomembrane hybrid silica nanoparticles prepared from living and dead cell sources. “Raw264.7” represents cell lysates. “Living cells” and “Dead cells” represent biomembrane hybrid silica nanoparticles prepared with iodixanol and sucrose, respectively. (A) Full coverage evaluation using fluorescence quenching of NBD by dithionite. (B) Overlap of proteins identified from each sample. (C) Graphical representation of Spearman’s rank correlation matrix. Heat-map shows the Spearman’s rank correlation between samples. Each column and row define individual technical replicates. In the heat-map, colors represent the *P*-values of Spearman correlations. (D) Volcano plot showing the −log_10_*P*-values as a function of the log_2_ ratios between nanoparticles prepared with living cells and dead cells. Horizontal and vertical axes represent the log_2_ ratio of the fold change and −log_10_ (Welch’s t-test, P value), respectively. Distribution of cells of proteins enriched in each of the particles prepared with (E) living cells and (F) dead cells.

Biomembrane hybridized nanoparticles prepared using living cells as a source have a high rate of complete coverage, which may result in an efficient coating of biomembrane-derived substances on the particle surface. To clarify that the prepared biomembrane hybridized nanoparticles inherit the membrane proteins of the source cells, SDS pages (sodium dodecyl sulfate-polyacrylamide gel electrophoresis) were performed on the particles and the extracted cell membranes (Figure S2A). No significant differences were observed between the cell membranes and the particles. We also performed western blotting for CD36 and CD44, which were reported as abundant proteins in Raw264.7 cell membranes (Figure S2B). These marker proteins were detected in the extracted plasma membrane and in the prepared particles. These results indicated that the proteins on the cell membrane were modified on the surface of the particles. Thus, the prepared nanoparticle could be nanocarriers with the characteristics of Raw264.7 cell membranes.

The biological conditions of the source cell could greatly influence the protein components on the prepared nanoparticles. Then, we analyzed the proteome profiling of the biomembrane hybridized nanoparticles without any bias and compared the results of the cell lysates and each nanoparticle. To this end, the digested peptides from the nanoparticles and the original cell lysate were prepared and compared with their proteome profiles. To confirm reproducibility, the LC-MS/MS experiments were repeated three times with three aliquots. Venn diagrams compared the lists of identified proteins from the original cells and the prepared nanoparticles (Figure 3B). Here, unique proteins identified in at least two of the three trials were used to increase the reliability, and 2632 proteins were commonly identified. According to gene ontology (GO) annotation obtained from the UniProt database using the David 6.8 database, 231 proteins identified only in the biomembrane hybridized particles prepared with living cells were assigned the “membrane” related keyword as cellular components (Figure S3), although 39 proteins nanoparticles prepared with dead cells and 101 proteins of cell lysate were not assigned these keywords^41^. On the other hand, the majority of proteins identified only in hybrid nanoparticles prepared with dead cells using sucrose or cell lysate were “intracellular” related keywords, including “cytoplasm” (Figure S3). We then calculated Spearman’s rank correlation between EV samples with the label-free quantification (LFQ) intensities, which was based on precursor signal intensity (Figure 3C). The correlation coefficients showed that each protein expression level was best correlated with itself across the three trials. Remarkably, there was a sufficient difference in protein composition between the particles prepared with living cells and other samples, indicating that specific cellular components were extracted by the cell layer penetration process. To quantitively investigate the upregulated proteins in the hybrid nanoparticle prepared with living cells and dead cells, the fold change values and *P*-values were calculated using LFQ intensities by Welch’s t-test. A fold change > 1 and *P*-value <0.05 were set as the criteria to select upregulated proteins. Nanoparticles prepared with living cells and dead cells contained 488 proteins and 672 proteins upregulated, respectively. They were depicted in volcano plots (Figure 3D). According to GO annotation, the “membrane” related keyword was annotated as the most over-represented GO terms in nanoparticles prepared with living cells and “intracellular” related keywords was annotated with dead cells (Figure S4). These tendencies were also observed with the same comparison with the cell (Figure S5). Briefly, the original cellular components of the enriched proteins were clearly different depending on the source cell conditions, and our method with living cells could emphasize the properties of membranous proteins on nanoparticle surfaces. (Figure 3E, 3F).

### Biomembrane hybridized nanoparticles reflect source cell membrane characteristics

Proteome profiling demonstrated that Biomembrane hybridized nanoparticles prepared with living cells could have the potential as superior DDS carriers, receiving the benefit of the membranous proteins. Characterizing the heterogeneous distribution of the prepared nanoparticle could be essential for quality control of biomedical applications, we investigated the molecular properties of the nanoparticles at the single-particle level. Here, we used imaging flow cytometry to evaluate the presence of lipid bilayer coatings by co-localization of fluorescence. Firstly, biomembrane hybridized nanoparticles were prepared using fluorescently stained cells for single-particle analysis. A 60x lens was used in the measurements with the following measurement conditions: flow rate as slow as possible, laser at 488 nm and 642 nm at maximum power. First, flow control beads were distinguished, and particles with confirmed FITC fluorescence were defined as silica nanoparticles (Figure S6, 4A). In the measurement, fluorescence images were acquired for each sample of 10,000 particles. When bare silica nanoparticles were measured using the prepared gating, no fluorescence of the cell membrane staining agent was detected. The prepared hybrid nanoparticles were then measured (Figure 4B). The results showed that fluorescence from the silica nanoparticles and the cell membrane staining agent were co-localized in the hybrid nanoparticles. This suggested that the silica nanoparticles are complex with biomembrane components. The fluorescence of the membrane staining agent was detected from 91% of the silica nanoparticles. This result suggested that almost all of the added silica nanoparticles were complexed with membrane components by using our method. In brief, compared to conventional coating methods, our method was achieved to complex nanoparticles and membrane components with high efficiency.

**Figure 4.**
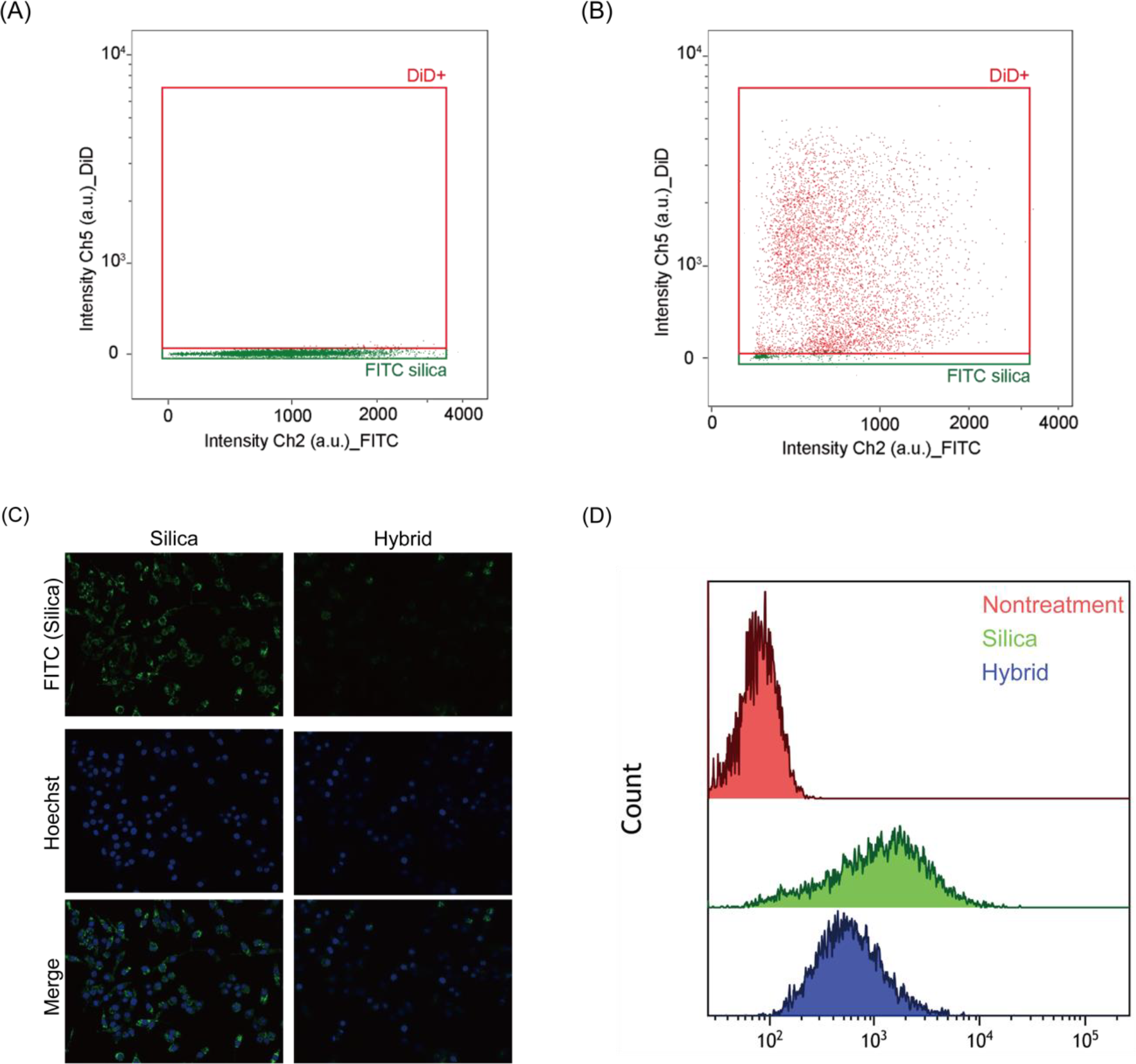
Evaluation of biomembrane hybrid silica nanoparticles prepared from live cell sources as drug delivery carriers. Results of single-particle analysis of bare (A) silica nanoparticles and (A)biomembrane hybridized nanoparticles using imaging flow cytometry. (C) Uptake behavior of bare silica nanoparticles and biomembrane hybrid nanoparticles by Raw264.7 cells. (D) Statistical evaluation of the uptake of each particle by Raw264.7 cells by flow cytometry.

Macrophage membranes could offer the nanoparticles the ability to circumvent macrophage clearance^42^. In the context of an exclusive biological system, this is a crucial concept for avoiding the mononuclear phagocyte system and delivering nanomaterials to target cells. We then investigated the uptake behavior of the hybridized nanoparticles by macrophages (Figure 4C). After incubating the macrophages with FITC-stained bare silica nanoparticles, strong fluorescence was observed in the cells, suggesting that macrophages efficiently uptake the nanoparticles by phagocytosis (Figure 4C; the left). On the other hand, FITC fluorescence was hardly observed in the cells after incubating the cells with biomembrane hybridized silica nanoparticles prepared with living macrophages (Figure 4C; the right). Statistical analysis of the uptake of silica nanoparticles using flow cytometry also suggested that uptake was significantly reduced with biomembrane hybridized silica nanoparticles (Figure 4D). In HeLa cells, the coating of the living biomembrane increased the uptake efficiency. This suggested that target specificity was enhanced by surface receptors inherited from the living cell membrane (Figure S7). These results suggested that the efficient coating of the macrophage membrane could give the particles the properties of the membrane and the ability to evade clearance. Overall, our method enabled us to coat nanomaterials effectively with membranous cellular components from living cells and provide a potential targeted delivery platform reflecting the functionality of the membrane.

In summary, we proposed a novel methodology to coat membrane components onto nanoparticle surfaces by forming cell layers at solution interfaces of varying densities. We showed that this approach could be used for complex membrane components with particles directly from living cells in a cell-nondestructive manner. In the evaluation, a comparison of hybrid nanoparticles prepared from living and dead cells as a source highlighted the advantage of using living cells as a source. In order to provide the nanoparticles with an effective biointerface, the integrity of the membrane is important^43^. In contrast to the case with dead cells, complete coverage was improved when live cells were used as the source. It should also be emphasized that the complete coverage rate is about 20%, which is higher than that of conventional coating methods. Furthermore, statistical single-particle analysis of the nanoparticles using imaging flow cytometry revealed that almost all of the silica nanoparticles that penetrated the cells with this method were complexed with the biomembrane. The absence of bare particles in the obtained particles suggested that the coating efficiency of this method was high. This suggested that the condition of the source membrane could have a significant effect on the coating rate and should motivate further studies on how to achieve complete cell membrane coating.

## Conclusion

In this study, we also performed a comprehensive analysis of proteins complexed with particles prepared from living and dead cells, and comprehensive and statical proteome profiling revealed that the use of living cells as a source significantly extracted intracellular membrane components, in contrast to particles prepared from dead cells or cell lysates. This was a surprising result and indicated that the method could efficiently prepare biomembrane hybridized nanoparticles without prior extraction of membrane components. Few studies have mentioned the comprehensive analysis of proteins contained in biomembrane hybridized nanoparticles, which may serve as a new indicator to characterize biomembrane hybridized nanoparticles. Future analysis of the proteins extracted when various types of particles are added to layers of cells of various types and states will lead to the construction of effective biointerfaces. Nanoparticles that are efficiently coated with the source cell membrane are expected to acquire the complex functions of the membrane. The biomembrane hybridized nanoparticles prepared from macrophages were complexed with characteristic membrane antigens and acquired the ability to evade macrophage clearance. In this study, membranes were coated on the nanoparticles using living cells as a source. Therefore, compared to previous membrane coatings, the proteins complexed may be different, and these differences may affect the function of the coated membrane. In the future, we expect to be able to design carriers that take advantage of the complex membrane functions predicted from the extracted proteins.

## Materials

HEPES, KOH, Sucrose (Ultra Pure), Succinic Anhydride, Magnesium sulphate, Hydrochloric Acid, SDS sample buffer, Dithiothreitol and Ethanol (99.5%), 4% Paraformaldehyde Phosphate Buffer Solution were purchased from FUJIFILM Wako Pure Chemical Corp (Osaka, Japan). Green fluorescent silica particles of 100 nm diameter sicastar-greenF and sicastar-greenF(plain, 100 nm) were purchased from micromod Partikeltechnologie GmbH (Roctock, Germany). Optiprep was purchased from COSMO BIO CO., LTD (Tokyo, Japan). (3-Aminopropyl)triethoxysilane and Trizma base were purchased from Merck (CA, USA). NBD-X was purchased from Invitrogen (MA, USA). CellBrite(R) Red Cytoplasmic Membrane Dye was purchased from Biotium (CA, USA). PBS (pH 7.4) and DMEM, high glucose were purchased from Thermo Fisher Scientific (MA, USA). RIPA Buffer(10x), Bullet Blocking One and Bullet CBB Stain One were purchased from NACALAI TESQUE, INC. (Kyoto, Japan). 10x/ Tris/Glycine/SDS buffer was purchased from BIO-RAD (CA, USA). CD36 Polyclonal antibody (18836-1-AP) and CD44 Monoclonal antibody (60224-1-Ig) were purchased from proteintech (IL, USA). m-IgG? BP-HRP was purchased from Santa Cruz Biotechnology (TX, USA). Goat Anti-Rabbit IgG H&L (HRP) preadsorbed was purchased from abcam (Cambridge, UK). ECL Western Blotting Detection Reagent was purchased from GE Healthcare (CHI, USA). Hoechst 33342 solution was purchased from DOJINDO (Kumamoto, Japan). Trypan Blue Stain was purchased from Logos biosystems (Gyeonggi-do, Korea)

### Cell culture

Raw264.7 cells were cultured in DMEM containing 10% FBS (Thermo Scientific, MA, USA) and 1% Gibco Antibiotic-Antimycotic (AA) (Thermo Scientific, MA, USA).

### Cell layer formation and penetration of nanoparticles

In a 3PC tube (Eppendorf Himac Technologies Co., Ltd, Ibaraki, Japan), Raw264.7 cells (1×107 cells) suspended in 500 µL of 60 wt% sucrose solution and 2 mL of HEPSE/KOH buffer (pH=7.4, 50 mM) were stacked. The cells were layered by ultracentrifugation (197,000 g, 5 min, 4°C). 0.1 mg of silica nanoparticles were added and ultracentrifuged again (197000 g, 2 h, 4°C) to allow the silica nanoparticles to penetrate the cell layers. The supernatant (2.3 mL) was removed and purified by centrifuge (20000 g, 10 min, 4°C) and sonication (28 kHz, 5 min). Cell viability after ultracentrifugation was also determined by trypan blue staining. When Iodixanol was used, a 30 wt% iodixanol solution was used to suspend the cells, and centrifugation after particle addition was changed to 197000 g, 10 min, 4°C.

### Evaluation of biomembrane hybrid nanoparticles

The particle size and Z voltage of the prepared particles were measured using a Zetasizer Nano ZS (Malvern Instruments, Worcestershire, UK). Fluorescence from silica nanoparticles was measured using an FP-6500 (JASCO, Tokyo, Japan). Protein concentrations were also determined by the Pierce™ BCA Protein Assay.

### Calculation of coverage by fluorescence quenching

NBD-modified silica nanoparticles were also used to prepare biomembrane hybrid nanoparticles for the coverage evaluation. 5 mg of silica nanoparticles were dispersed in 10 mL of ethanol. Then 20 µL of (3-Aminopropyl)triethoxysilane was added and reacted for 2 hours. The solution was centrifuged at 20000 × g for 10 min. The supernatant was removed, and the same volume of ethanol was added two times. 2 mg of NBD-X was added and reacted overnight. The mixture was washed twice and stirred for 1 hour. 9 mg of succinic anhydride was added, stirred for 5 hours, washed three times, and dispersed in HEPES buffer.

A 100 µL solution of each nanoparticle (NBD labeled, 500 µg/mL) was added to the fluorometer cell and time-resolved measurements were performed on an FP-6500 (JASCO, Tokyo, Japan). Measurements were taken at 10 second intervals, and at 120 seconds, 20 µL of 0.1 M sodium dithionite was added to quench the fluorescence. Normalized Fluorescence was calculated for analysis using the following equation. F_T_ is the total fluorescence of the sample before addition of dithionite, F_D_ is the fluorescence of the sample after addition of dithionite with dithionite and F_0_ is the background.

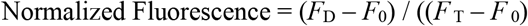

### Protein Digestion and Purification of Peptide Samples for LC-MS/MS Analysis

Samples were dried and digested using the phase-transfer surfactant (PTS)-aided trypsin digestion protocol as described previously^44^. Proteins were reduced with 10 mM dithiothreitol (DTT) (FUJIFILM Wako Pure Chemical Corporation, Osaka, Japan) for 30 min at 37 °C, followed by alkylation with 50 mM 2-iodoacetamide (IAA) (FUJIFILM Wako Pure Chemical Corporation) for 30 min at room temperature in the dark. The samples were diluted to 2 M urea with 50 mM ammonium bicarbonate. The proteins were digested with 1 μg of lysyl endopeptidase (LysC) (FUJIFILM Wako Pure Chemical Corporation) and 1 μg of trypsin (Promega, Tokyo, Japan) overnight at 37 °C in a shaking incubator. The resulting peptides were acidified with 0.5% trifluoroacetic acid (TFA, final concentration) and fractionated with a StageTip containing SDB-XC (upper) and SCX (bottom) Empore disk membranes (GL Sciences, Tokyo, Japan). Peptides were washed with 0.1% TFA with 5% ACN and 0.1% TFA with 80% ACN. Then they were eluted from the tip by 4% TFA with 30% ACN in 500 mM ammonium acetate and 30% ACN in 500 mM ammonium acetate. The sample solution was evaporated in a SpeedVac (Thermo Fisher Scientific), and the residue was resuspended in 0.5% TFA with 5% ACN. Finally, the peptides were desalted again by StageTip with SDB-XC Empore disk membranes and suspended in a loading buffer (0.5% TFA with 5% ACN) for subsequent LC-MS/MS analyses. After digestion, peptide concentration was measured on NanoDrop (Thermo Fisher Scientific) with an absorbance at 205 nm and an extinction coefficient of 31 and each peptide sample concentration was adjusted to 160 ng/μL in 0.5% TFA with 4% ACN^45^.

### LC-MS/MS Analysis

NanoLC-MS/MS analyses were performed using a Q-Exactive instrument (Thermo Fisher Scientific), which was connected to an UltiMate 3000 pump (Thermo Fisher Scientific) and an HTC-PAL autosampler (CTC Analytics). Peptides were separated on pulled-in-house needle columns (150 mm length, 100 μm inner diameter, 6 μm needle opening) packed with ReproSil-Pur 120 C18-AQ 3 μm RP material (Dr. Maisch, Ammerbuch, Germany)^46^. The samples were applied by 5 μL of full loop injection, and the flow rate was 500 nL/min. Separation was achieved by using a four-step linear gradient of 4 to 10% ACN in 5 min, 10 to 40% MeCN in 60 min, 40 to 99% MeCN in 10 min, and 99% MeCN for 10 min with 0.5% TFA. The electrospray voltage was set to 2.4 kV in positive mode. The full MS scan was acquired with a mass range of 350–1500 m/z, a resolution of 70,000, an automatic gain control (AGC) target of 3e, and a maximum injection time of 100 ms. The MS/MS scan was performed by the Top10 method with a resolution of 17,500, the AGC target of 1e, a maximum injection time of 100 ms, and an isolation window of 2.0 Th. The precursor ions were fragmented by higher-energy collisional dissociation, with a normalized collision energy of 27%.

### Database Searching

For all experiments, the raw MS data files were analyzed by MaxQuant v2.0.3.0^47^. Peptides and proteins were identified by an automated database searching using Andromeda against the SwissProt Database of mus musculus (mouse) (version 2023-08, 17,784 protein entries) with a precursor mass tolerance of 20 ppm for the first search, 4.5 ppm for the main search, and a fragment ion mass tolerance of 20 ppm. The enzyme was set as trypsin/P, with two missed cleavages. Cysteine carbamidomethylation was set as a fixed modification. Methionine oxidation and acetylation on the protein N-terminus were set as variable modifications. The search results were filtered with FDR < 1% at the peptide spectrum match (PSM) and protein levels. The match-between-run algorithm (MBR) was utilized through the “Identification” subtab in the “Global Parameters” tab of MaxQuant to mitigate the missing value problem. The default settings for MBR were used (0.7 min match window and 20 min alignment time). Proteins that have “only identified by site”, “potential contaminants”, and “reverse sequences” were removed for data analysis.

### Single particle analysis

For single-particle analysis, biomembrane hybrid nanoparticles were prepared using cells whose membranes had been labeled fluorescently with CellBrite. The measurements were performed on an AMNIS ImageStreamX Mark II Flow Cytometer (Luminex, TX, USA). Calibration was performed before each sample measurement. The flow velocity was calibrated from the average velocity of SpeedBeads (Luminex, TX, USA) flowing in the channel. For measurements of biomembrane hybrid nanoparticles, the signals were detected at Ch4 for brightfield, Ch2 for FITC-modified silica nanoparticles, Ch5 for CellBright in the cell membrane, and Ch6 for SSC. The same gating was used for all samples. Samples were measured for 10,000 particles detected in the region of silica nanoparticles. Analysis of the fluorescence intensity of each object was performed by IDEAS (Luminex, TX, USA).

### Evaluation of hybridization of cell membrane antigens with nanoparticles

Cell membranes derived from Raw264.7 were prepared as follows: Raw264.7 cells were suspended in ice-cold Tris-magnesium buffer (2.5 × 10^7^ cells/mL) and passed through an extruder (without polycarbonate membrane) 20 times to disrupt the cells. 1 M sucrose solution was mixed was mixed to a final concentration of 0.25 M and centrifuged (2000 g, 4 °C, 10 min). The supernatant was collected and further centrifuged (3000 g, 4 °C, 30 min). The resulting cell membrane was washed with ice-cold 0.25 M sucrose solution and isolated by centrifugation (3000 g, 4 °C, 30 min).

Samples were mixed with 10 μL of SDS sample buffer containing DTT and heated (70°C, 15 min) for reduction. SDS-PAGE was then performed using 7.5% e-PAGEL. The gels were then stained.

For Western blotting, the gel bands were transferred to PVDF after performing SDS-PAGE. To detect membrane antigens, the membrane was incubated overnight at 4°C with a 2000-fold dilution of the primary antibody. The membrane was then incubated with diluted secondary antibodies (CD36: 1:1000, CD44: 1:2000) for one hour. Membranes were stained with ECL Western Blotting Detection Reagent and visualized using LAS-4000 EPUV mini (FUJIFILM, Tokyo, Japan).

### Cellular uptake of nanoparticles

Raw264.7 cells (40,000 cells/well) were seeded in a multi-well bottom dish. 2 µg of each silica nanoparticles and biomembrane hybrid nanoparticles was added to cells and incubated at 37°C. After each addition, cells were fixed with 4% paraformaldehyde and observed under an BZ-X800 confocal (Keyence, Tokyo, Japan).

For analysis by flow cytometry, 100000 cells/well Raw264.7 cells were first seeded in 12 wells. Silica nanoparticles and biomembrane hybrid nanoparticles were added to cells for 20 µg each and incubated at 37°C for 4 hours. Intracellular FITC fluorescence was then measured by BD LSRFortessa™ Cell Analyzer (BD bioscience, NJ, USA). Analysis was performed on Kluza (Beckman Coulter Life Sciences, CA, USA).

### Statistical Analysis

All measurements were performed using at least three independent samples. Comparisons of the particle size and ζvoltage were evaluated using Welch’s t test. P-values less than 0.05 were considered statistically significant. In addition, statistical significance is marked in figures with *** for p < 0.005, ** for p < 0.01 and * for p < 0.05. All statistical analyses were performed using Graphpad Prism 9 (GraphPad Software, Inc., CA, USA).

## Data Availability

The MS raw data and analysis files have been deposited at the ProteomeXchange Consortium (http://proteomecentral.proteomexchange.org) via the jPOST partner repository (https://jpostdb.org) with the data set identifier JPST002365^48^.

## Acknowledgment

This work was supported by grants from the Japan Society for the Promotion of Science (JSPS) Grants-in-Aid for Scientific Research: Nos. JP16H06313, JP22H00585 (KA), JP16H03842, JP21K18324, JP22H02199, JP21H04954 (YS), JP23K13774 and 21K14652 (EK).

## TOC

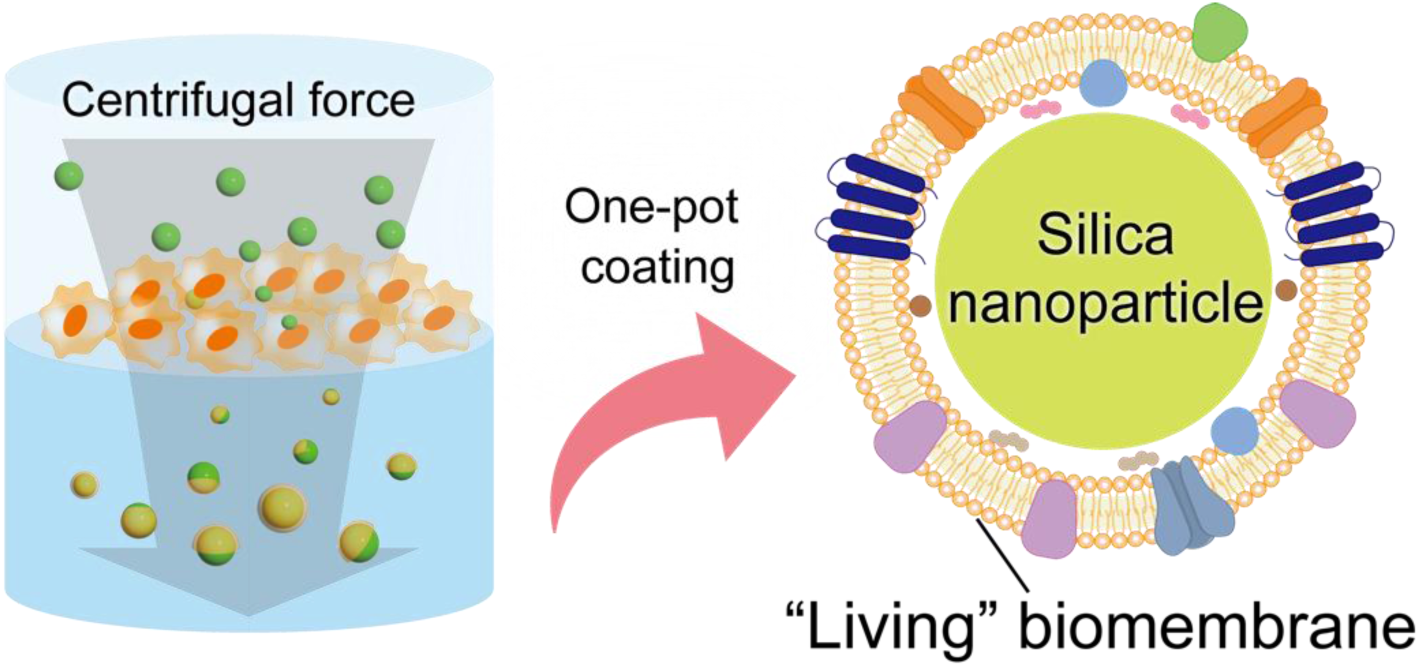

In this paper, silica nanoparticles were efficiently hybridized with living cell membrane by passing them through cell layers prepared by density gradient centrifugation. As a result of comprehensive proteome profiling, biomembrane hybridized nanoparticles mainly included membranous proteins originating from cell membranes and organelle membranes on the surface. Furthermore, the coated cell membrane effectively suppressed the phagocytic activity of Raw264.7 cells.

## References

1. Mitchell MJ, Billingsley MM, Haley RM, Wechsler ME, Peppas NA, Langer R. Engineering precision nanoparticles for drug delivery. Nat. Rev. Drug Discov. 20, 101–124 (2021).

2. Yetisgin AA, Cetinel S, Zuvin M, Kosar A, Kutlu O. Therapeutic Nanoparticles and Their Targeted Delivery Applications. Molecules 25, (2020).

3. Kim D, Kim J, Park YI, Lee N, Hyeon T. Recent Development of Inorganic Nanoparticles for Biomedical Imaging. ACS Cent. Sci. 4, 324–336 (2018).

4. Waheed S, Li Z, Zhang F, Chiarini A, Armato U, Wu J. Engineering nano-drug biointerface to overcome biological barriers toward precision drug delivery. J. Nanobiotechnology 20, 395 (2022).

5. Shi L, Zhang J, Zhao M, Tang S, Cheng X, Zhang W, Li W, Liu X, Peng H, Wang Q. Effects of polyethylene glycol on the surface of nanoparticles for targeted drug delivery. Nanoscale 13, 10748–10764 (2021).

6. Zhang W, Taheri-Ledari R, Ganjali F, Mirmohammadi SS, Qazi SF, Saeidirad M, KashtiAray A, Zarei-Shokat S, Tian Y, Maleki A. Effects of morphology and size of nanoscale drug carriers on cellular uptake and internalization process: a review. RSC Adv. 13, 80–114 (2023).

7. Jiang L, Lee HW, Loo SCJ. Therapeutic lipid-coated hybrid nanoparticles against bacterial infections. RSC Advances 10, 8497–8517 (2020).

8. Stater EP, Sonay AY, Hart C, Grimm J. The ancillary effects of nanoparticles and their implications for nanomedicine. Nat. Nanotechnol. 16, 1180–1194 (2021).

9. van Niel G, Carter DRF, Clayton A, Lambert DW, Raposo G, Vader P. Challenges and directions in studying cell–cell communication by extracellular vesicles. Nat. Rev. Mol. Cell Biol. 23, 369–382 (2022).

10. Song B, Sheng X, Justice LJ, Lum KK, Metzger JP, Cook CK, Kotas CJ, Cristea MI. Intercellular communication within the virus microenvironment affects the susceptibility of cells to secondary viral infections. Sci. Adv. 9, eadg3433 (2023).

11. Herrmann IK, Wood MJA, Fuhrmann G. Extracellular vesicles as a next-generation drug delivery platform. Nat Nanotechnol. 16, 748–759 (2021).

12. Kalluri R, LeBleu VS. The biology, function, and biomedical applications of exosomes. Science 367, eaau6977 (2020).

13. Gurung S, Perocheau D, Touramanidou L, Baruteau J. The exosome journey: from biogenesis to uptake and intracellular signalling. Cell Commun. Signal. 19, 47 (2021).

14. Mathieu M, Martin-Jaular L, Lavieu G, Théry C. Specificities of secretion and uptake of exosomes and other extracellular vesicles for cell-to-cell communication. Nat. Cell Biol. 21, 9–17 (2019).

15. Zöller M. Tetraspanins: push and pull in suppressing and promoting metastasis. Nat. Rev. Cancer 9, 40–55 (2009).

16. Shimoda A, Miura R, Tateno H, Seo N, Shiku H, Sawada S, Sasaki Y, Akiyoshi K. Assessment of Surface Glycan Diversity on Extracellular Vesicles by Lectin Microarray and Glycoengineering Strategies for Drug Delivery Applications. Small Methods 6, 2100785 (2022).

17. Hoshino A, Costa-Silva B, Shen TL, Rodrigues G, Hashimoto A, Tesic MM, Molina H, Kohsaka S, Di Giannatale A, Ceder S, Singh S, Williams C, Soplop N, Uryu K, Pharmer L, King T, Bojmar L, Davies EA, Ararso Y, Zhang T, Zhang H, Hernandez J, Weiss MJ, Dumont-Cole DV, Kramer K, Wexler HL, Narendran S, Schwartz KG, Healey HJ, Sandstrom P, Labori JK, Kure HE, Grandgenett MP, Hollingsworth AM, Sousa M, Kaur S, Jain M, Mallya K, Batra KS, Jarnagin RW, Brady SM, Fodstad O, Muller V, Pantel K, Minn JA, Bissell JM, Garcia AB, Kang Y, Rajasekhar KV, Ghajar MC, Matei I, Peinado H, Bromberg J, Lyden D. Tumour exosome integrins determine organotropic metastasis. Nature 527, 329–335 (2015).

18. Más V, Melero JA. Entry of enveloped viruses into host cells: membrane fusion. Subcell. Biochem. 68, 467–487 (2013).

19. Chazal N, Gerlier D. Virus entry, assembly, budding, and membrane rafts. Microbiol. Mol. Biol. Rev. 67, 226–237, (2003).

20. O’Brien K, Breyne K, Ughetto S, Laurent LC, Breakefield XO. RNA delivery by extracellular vesicles in mammalian cells and its applications. Nat. Rev. Mol. Cell Biol. 21, 585–606 (2020).

21. Dimitrov DS. Virus entry: molecular mechanisms and biomedical applications. Nat. Rev. Microbiol. 2, 109–122 (2004).

22. Alvarez-Erviti L, Seow Y, Yin H, Betts C, Lakhal S, Wood MJA. Delivery of siRNA to the mouse brain by systemic injection of targeted exosomes. Nat. Biotech. 29, 341–345 (2011).

23. Antimisiaris SG, Mourtas S, Marazioti A. Exosomes and Exosome-Inspired Vesicles for Targeted Drug Delivery. Pharmaceutics 10, (2018).

24. Charoenviriyakul C, Takahashi Y, Morishita M, Matsumoto A, Nishikawa M, Takakura Y. Cell type-specific and common characteristics of exosomes derived from mouse cell lines: Yield, physicochemical properties, and pharmacokinetics. Eur. J. Pharm. Sci. 96, 316–322 (2017).

25. Ferguson SW, Nguyen J. Exosomes as therapeutics: The implications of molecular composition and exosomal heterogeneity. J. Control. Release 228, 179–190 (2016).

26. Cheng G, Li W, Ha L, Han X, Hao S, Wan Y, Wang Z, Dong F, Zou X, Mao Y, Zhenget S. Self-Assembly of Extracellular Vesicle-like Metal–Organic Framework Nanoparticles for Protection and Intracellular Delivery of Biofunctional Proteins. J. Am. Chem. Soc. 140, 7282–7291 (2018).

27. Fang RH, Kroll AV, Gao W, Zhang L. Cell Membrane Coating Nanotechnology. Adv. Mater. 30, 1706759 (2018).

28. Fang RH, Gao W, Zhang L. Targeting drugs to tumours using cell membrane-coated nanoparticles. Nat. Rev. Clin. Oncol. 20, 33–48 (2023).

29. Dhas N, García CM, Kudarha R, Pandey A, Nikam NA, Gopalan D, Fernandes G, Soman S, Kulkarni S, Seetharam NR, Tiwari R, Wairkar S, Pardeshi C, Mutaliket S. Advancements in cell membrane camouflaged nanoparticles: A bioinspired platform for cancer therapy. J. Control. Release 346, 71–97 (2022).

30. Song M, Dong S, An X, Zhang W, Shen N, Li Y, Guo C, Liu C, Li X, Chen S. Erythrocyte-biomimetic nanosystems to improve antitumor effects of paclitaxel on epithelial cancers. J. Control. Release 345, 744–754 (2022).

31. Ma Y, Ma Y, Gao M, Han Z, Jiang W, Gu Y, Liu, Y. Platelet-Mimicking Therapeutic System for Noninvasive Mitigation of the Progression of Atherosclerotic Plaques. Adv. Sci. 8, 2004128 (2021).

32. Xiao T, He M, Xu F, Fan Y, Jia B, Shen M, Wang H, Shi X. Macrophage Membrane-Camouflaged Responsive Polymer Nanogels Enable Magnetic Resonance Imaging-Guided Chemotherapy/Chemodynamic Therapy of Orthotopic Glioma. ACS Nano 15, 20377–20390 (2021).

33. Shu X, Chen Y, Yan P, Shi Q, Yin T, Wang P, Liu L, Shuai X. Biomimetic nanoparticles for effective mild temperature photothermal therapy and multimodal imaging. J. Control. Release 347, 270–281 (2022).

34. Zou D, Wu Z, Yi X, Hui Y, Yang G, Liu Y, Tengjisi, Wang H, Brooks A, Wang H, Liu X, Xu PZ, Roberts SM, Gao H, Zhao C. Nanoparticle elasticity regulates the formation of cell membrane-coated nanoparticles and their nano-bio interactions. Proc. Natl. Acad. Sci. USA 120, e2214757120 (2023).

35. Qin M, Du G, Qiao N, Guo Z, Jiang M, He C, Bai S, He P, Xu Y, Wang H, Gong T, Zhang Z, Sun X. Whole-Cell-Mimicking Carrier-Free Nanovaccines Amplify Immune Responses Against Cancer and Bacterial Infection. Adv. Funct. Mater. 32, 2108917 (2022).

36. Liu Y, Luo J, Chen X, Liu W, Chen T. Cell Membrane Coating Technology: A Promising Strategy for Biomedical Applications. Nano-Micro Lett. 11, 100 (2019).

37. Liu L, Bai X, Martikainen M, Kårlund A, Roponen M, Xu W, Hu G, Tasciotti E, Lehto V. Cell membrane coating integrity affects the internalization mechanism of biomimetic nanoparticles. Nat. Commun. 12, 5726 (2021).

38. Mizuta R, Inoue F, Sasaki Y, Sawada S-i, Akiyoshi K. A Facile Method to Coat Nanoparticles with Lipid Bilayer Membrane: Hybrid Silica Nanoparticles Disguised as Biomembrane Vesicles by Particle Penetration of Concentrated Lipid Layers. Small 19, 2206153 (2023).

39. Lobb RJ, Becker M, Wen WS, Wong SFC, Wiegmans PA, Leimgruber A, Möller A. Optimized exosome isolation protocol for cell culture supernatant and human plasma. J. Extracell. Vesicles 4, 27031 (2015).

40. Kovacovicova K, Vinciguerra M. Isolation of senescent cells by iodixanol (OptiPrep) density gradient-based separation. Cell Prolif. 52, e12674 (2019).

41. Huang da W, Sherman BT, Lempicki RA. Systematic and integrative analysis of large gene lists using DAVID bioinformatics resources. Nat. Protoc. 4, 44–57 (2009).

42. Gao C, Huang Q, Liu C, Kwong HTC, Yue L, Wan J, Lee MYS, Wang R. Treatment of atherosclerosis by macrophage-biomimetic nanoparticles via targeted pharmacotherapy and sequestration of proinflammatory cytokines. Nat. Commun. 11, 2622 (2020).

43. Liu L, Liu L, Pan D, Chen S, Martikainen M, Kårlund A, Ke J, Pulkkinen H, Ruhanen H, Roponen M, Käkelä R, Xu W, Wang J, Lehto V. Systematic design of cell membrane coating to improve tumor targeting of nanoparticles. Nat. Commun. 13, 6181 (2022).

44. Masuda T, Tomita M, Ishihama Y. Phase Transfer Surfactant-Aided Trypsin Digestion for Membrane Proteome Analysis. J. Proteome Res. 7, 731–740 (2008).

45. Scopes RK. Measurement of protein by spectrophotometry at 205 nm. Anal. Biochem. 59, 277–282 (1974).

46. Ishihama Y, Rappsilber J, Andersen JS, Mann M. Microcolumns with self-assembled particle frits for proteomics. J. Chromatogr. A 979, 233–239 (2002).

47. Cox J, Mann M. MaxQuant enables high peptide identification rates, individualized p.p.b.-range mass accuracies and proteome-wide protein quantification. Nat. Biotechnol. 26, 1367–1372 (2008).

48. Okuda S, Watanabe Y, Moriya Y, Kawano S, Yamamoto T, Matsumoto M, Takami T, Kobayashi D, Araki N, Yoshizawa CA, Tabata T, Sugiyama N, Goto S, Ishihama Y. jPOSTrepo: an international standard data repository for proteomes. Nucleic Acids Research 45, D1107–D1111 (2016).

